# Intermittent Fasting Enhances Hippocampal Npy Expression to Promote Neurogenesis Following Traumatic Brain Injury

**DOI:** 10.1101/2021.04.13.439591

**Authors:** Shuqiang Cao, Manrui Li, Yuwen Sun, Wenjie Yang, Hao Dai, Yadong Guo, Yi Ye, Zheng Wang, Xiaoqi Xie, Xiameng Chen, Weibo Liang

**Author notes:** These authors contributed equally to this work (Shuqiang Cao, Manrui Li). Corresponding author: <u>Weibo Liang</u>, Department of Forensic Genetics, West China School of Basic Medical Sciences and Forensic Medicine, Sichuan University, Chengdu, China. <u>Xiameng Chen</u>, Department of Forensic Pathology and Forensic Clinical Medicine, West China School of Basic Medical Sciences and Forensic Medicine, Sichuan University, Chengdu, China.

## Abstract

Interventions for preventing cognitive dysfunction post traumatic brain injury (TBI) is limited. Given that adult hippocampal neurogenesis (AHN) after brain injury contributes to cognitive recovery, and that the AHN is potentially affected by nutritional factors, we asked whether fasting could promote AHN and thus ameliorates cognitive defects after TBI. Here we show that a one-month intermittent fasting (IF) regimen enhanced proliferation of neural stem cells (NSCs) in the subgranular zone (SGZ) of hippocampus 3 days post TBI, as well as improved cognitive performance in Morris water maze (MWM) test. Furthermore, an increase in hippocampal Npy expression was detected in IF group after injury, compared to the mice fed ad libitum (AL), and locally knock-down of Npy in vivo attenuated the aforementioned effects of IF in TBI. These findings suggest that IF promotes AHN following TBI by a mechanism involving enhancement of Npy expression, which may offer novel interventions that might prevent cognitive dysfunction caused by injury.

## Introduction

The link between TBI and long-term cognitive dysfunction is increasingly established[1, 2], while therapeutic interventions to prevent this impairment is quite limited, resulting in an enormous social and economic burden.

It is known that the hippocampus is critical in maintaining cognitive function, especially in learning and memory storage[3]. Hippocampal dysfunction-a major aspect of pathology following TBI, usually acts as the cause of cognitive impairment. Evidences have repeatedly shown a relationship between adult hippocampal neurogenesis (AHN) and its dependent cognitive function in rodents[4]. Impairments in the use of spatial strategies and retention of spatial locations in the water maze tests have been found under AHN ablation via administration of methylazoxymethanol acetate (MAM) or TMZ100[5–7]. Subsequently, defects in spatial memory and electrophysiological properties of the dentate gyrus (DG) were confirmed under genetic ablation of AHN[8, 9], highlighting the role of AHN in cognitive function. In addition, in an animal study involving TBI, the ability to learn spatial memory tasks was also found lost in mice treated with temporally specified ablation of neurogenesis[10]. To this end, targeting AHN may provide novel therapeutic options for the treatment or prevention of cognitive defects after brain injury.

Recently, adult neurogenesis has been found to be affected by nutritional factors[11]. Also, for the past few years, positive relationship between food restriction and hippocampus-dependent cognition has been proposed[12, 13]. Intermittent fasting (IF) is a novel strategy of energy restriction, which entails drastic reductions in food intake for varying periods of time. Recent efforts have shown that cognition in multiple domains could be enhanced by IF[14–17]. In clinic trials, long-term energy restriction was demonstrated to induce improvements in executive function, verbal memory and global cognition in adults with overweight and mild cognitive impairments[18]. Also, another clinical study involving older adults revealed the relationship between a short-term food restriction and an improvement in verbal memory[19]. Hence, there is a great possibility that IF may enhance injury-induced AHN, improving its related cognitive performance in TBI.

This study aimed to explore the impact of IF on the AHN and the hippocampus-dependent cognitive recovery process responding to moderate TBI, as well as the underlying mechanisms. Gaining this detailed understanding of the link between IF and hippocampal behavior under TBI is paramount for the development of new interventions, alleviating the associated social, and economic burdens.

## Methods

### 1. Animals

8-10 weeks C57BL/6N mice, half male, half female, weighing from 21-23g were utilized in our study. The mice were randomly assigned to each group, with 10~18 mice per group. All the mice were housed in the AAALAC (Association for Assessment and Accreditation of Laboratory Animal Care International) approved animal facility under a 12-h light/dark cycle, 22–25 °C, 50% relative humidity. Animal procedures were performed based on the Ethical Research Committee guidelines of Sichuan University of Medical Sciences.

### 2. IF

We used an alternate day fasting regimen in this study. Mice were fasted one day and had free access to food and water the next day. On fasting day, food pillets were removed from the cage, while water was available. The control group was fed on a standard mice diet ad libitum.

### 3. CCI Model

Briefly, mice were anesthetized with 4% isoflurane and maintained on a 2% concentration. Then the mice were placed and fixed on a stereotaxic frame. A midsagittal incision was made in the scalp and periosteum under sterile conditions. A 3.5mm diameter craniotomy was made posterial to the coronal suture with a surgical drill, and then a 3mm flat hitting tip was vertically placed on the exposed dural surface for cortical impact. After the impact, bone widow was covered by sterile plastic film, and skin was intermittent sutured and disinfected. For more details, please see[20].

### 4. Western Blot Assay, H&E staining, Immunohistochemistry (IHC) and qPCR

As previously described[20–22].

### 5. Morris Water Maze (MWM) Assay

The experiment lasted for 5 days and all mice were scheduled to be trained 4 times per day for a fixed period of time. During training, the mice were placed into the pool through four entry points. The time that mice took to enter the water to find the submerged hidden platform and stand on it was recorded as latency. If the mice could find it themselves, they were left on the platform. If the mice did not find the platform within 90 s, they were gently pulled on the platform for 10s. Each mouse was placed into the pool through four water inlet points with a 30s training interval. On the fifth day, the mice were placed in the water at the same inlet point for each quadrant, and the swimming path of the mice over 90 s was recorded. The time the mice stayed in the target quadrant platform was recorded and used to evaluate the spatial localization ability and long-term memory of the mice. The shorter latency and longer staying time indicate better spatial localization ability and long-term memory, which is associated with hippocampal-dependent learning ability. For more details, please see[20].

### 6. Intracerebroventricular (I.C.V.) injection

AAV-Npy-shRNA and AAV-scramble were packaged at a titer of 5.28×1010 IFU/mL and 12.30×1010 IFU/mL (purchased from Shanghai GenePharma Co., Ltd). Packaged Npy-shRNA and scramble were injected into I.C.V. according to the manufacturer’s instructions. For I.C.V, each anesthetized mouse received was placed and fixed in the stereotaxic frame and had midsagittal incision on the scalp. A 1.0- or 5.0-μl Hamilton syringe (Hamilton, Reno, NV) was used to inject specified quantities of AAV-Npy-shRNA and AAV-scramble in a volume of 0.5 μl into the right cerebral ventricle. (location: please see “results”).

### 7. Statistical analysis

All statistical analyses were performed and the figures were made using Graphpad Prism 8.4.2 software (San Diego, USA). Values are expressed as the mean ± standard deviation(SD). Differences between groups were analyzed using one-way or two-way ANOVA. A p value<0.05 was considered to be statistically significant.

## Results

### IF enhances AHN following TBI

Currently, the effect of IF on neurogenesis in TBI has not been reported. In this study, we used an alternate day fasting regimen (Fig1. A) based on a moderate controlled cortical impact (CCI) model to address this question. Histologically, we found that the 1 months IF resulted in an increase in the number of shuttle-shaped cells with more basophilic staining lining in SGZ (the neurogenic niche in hippocampus), morphologically alike the NSCs (Fig1. B). To validate the identity of these increased shuttle-shaped cells, we conducted immune fluorescence staining against the proliferative marker-ki67, as well as the NSC marker-sox2. As shown in Fig 2, ki67 positive cells in the SGZ was largely increased in the IF pre-conditioned group, compared to the AL fed group (Fig1. C), suggesting an effect of IF in increasing cell proliferation post TBI. Furthermore, almost all of the ki67 positive cells were co-stained with sox2, proving the NSC identity of these proliferating cells. Hence, our results suggested that the SGZ neurogenesis responding to TBI could be enhanced by long-term IF.

**Fig1.**
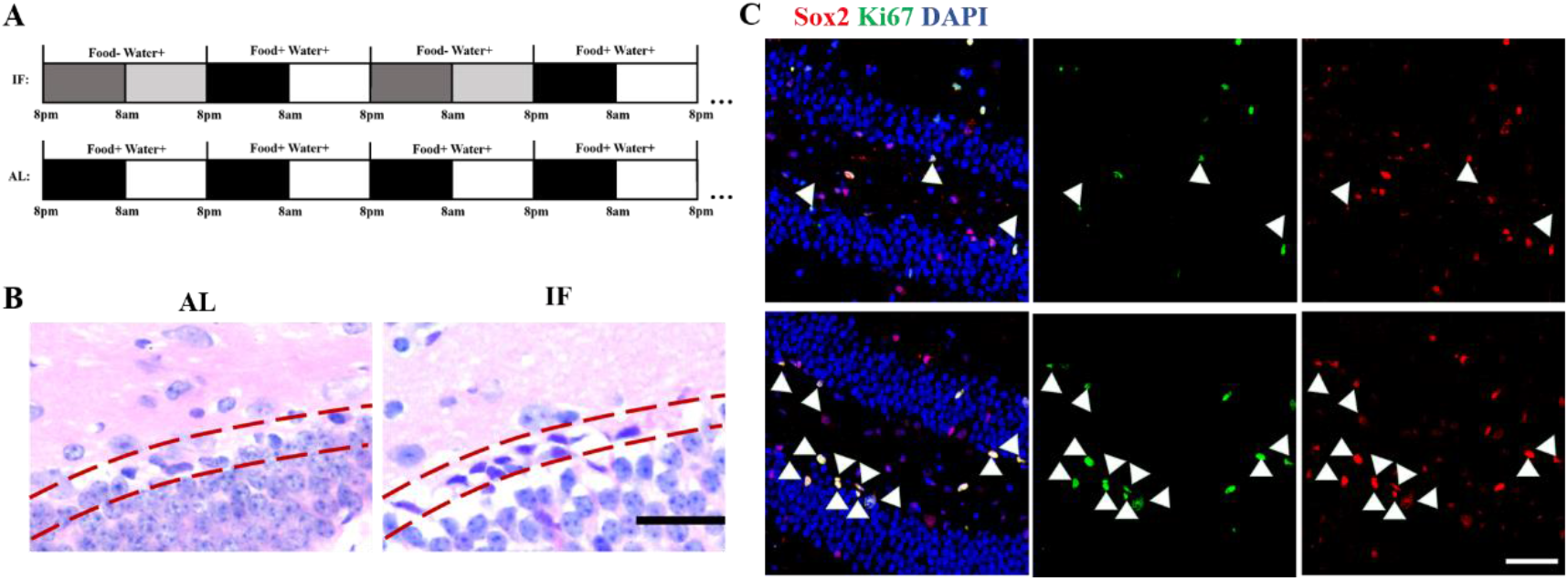
AHN following TBI was enhanced under IF pre-condition. A) alternate day fasting regimen. B) H&E staining showed the number of shuttle-shaped cells with basophilic staining was increased in the SGZ (indicated by dashed lines) in IF group on post injury day3, scale =50um. C) immune-staining showed that the ki67 and sox2 positive cells were increased in the SGZ in IF group on post injury day3, scale=100um.

**Fig2.**
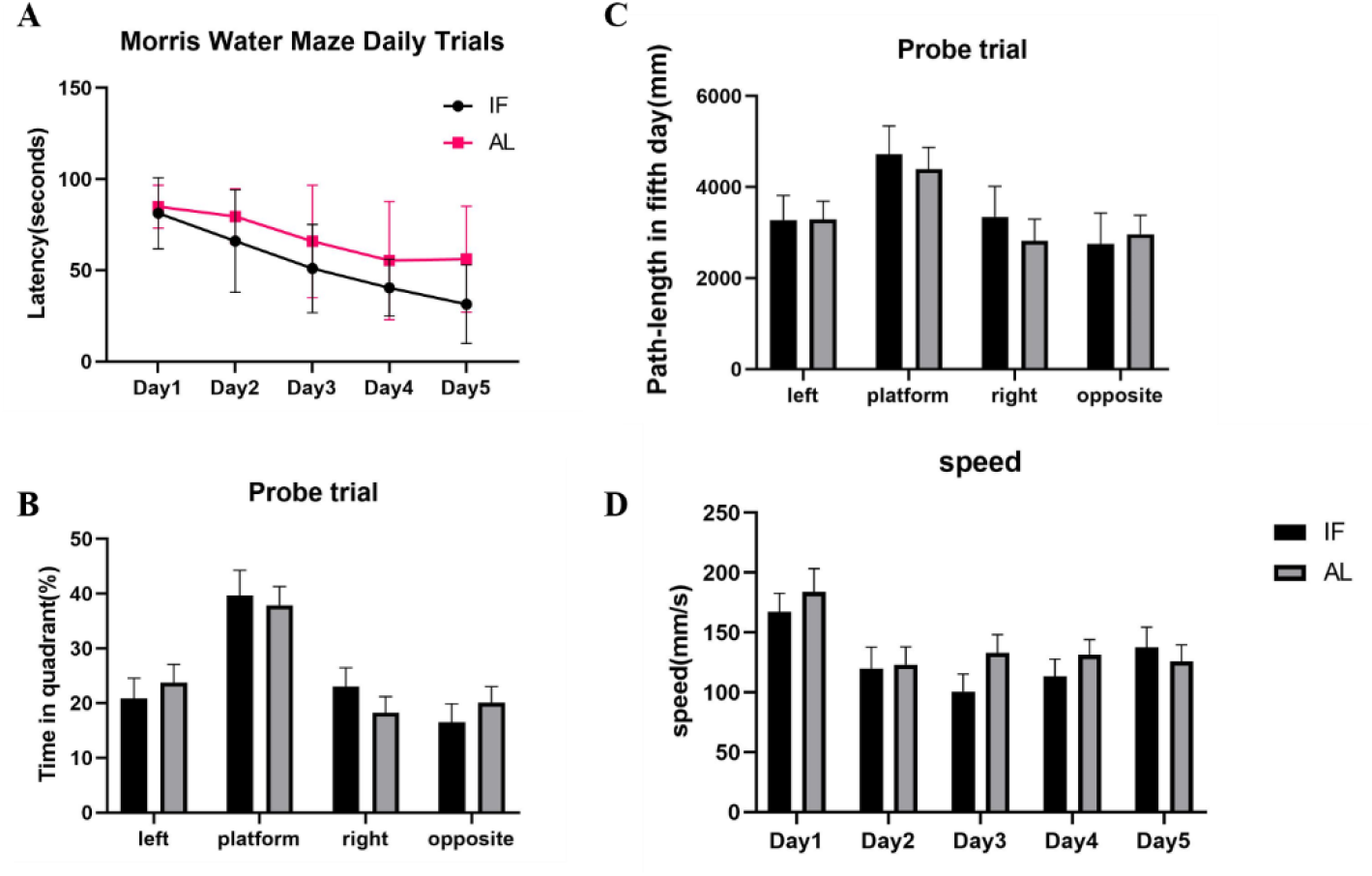
IF pre-treatment ameliorated cognitive dysfunction 1month post TBI. A) IF group exhibited significantly decreased goal latency in MWM test (P<0.05). B) time in quadrants C) path length and D) speed tests did not show significant differences between IF and AL groups.

### Cognitive dysfunction post TBI is ameliorated under IF

The hippocampal neurogenesis events are critical in cognition recovery following TBI. Hence, a de-facto standard for testing the hippocampal function-Morris water maze (MWM), was performed 1 month post injury to test the impact of IF on cognitive performance post injury (Fig2. A-D). Results showed that the IF preconditioning before a moderate TBI resulted in improved cognition post injury, as seen by significantly decreased goal latency (Fig2. A), demonstrating that the post injury cognitive defects are partly alleviated by IF pre-treatment.

### Enhanced Npy expression contributes to the effects of IF in TBI

To clarify the mechanisms underlying the neuroprotective effect of IF in TBI, we focused on genes involved in both neurogenesis promoting and food intake regulation. It has been known that a neuroprotective peptide-Npy, is largely increased in hypothalamus responding to fasting. Also, it has been suggested to participate in neurogenesis event. However, whether Npy in murine hippocampus could be induced by IF remains unclear. To address this, we conducted immuno-staining against Npy in IF and AL hippocampus, respectively. We found an obvious increase of Npy immunoactivity in IF hippocampus (Fig3. A). In addition, WB results further validated the elevated hippocampal Npy expression in the IF group (Fig3. B).

**Fig3.**
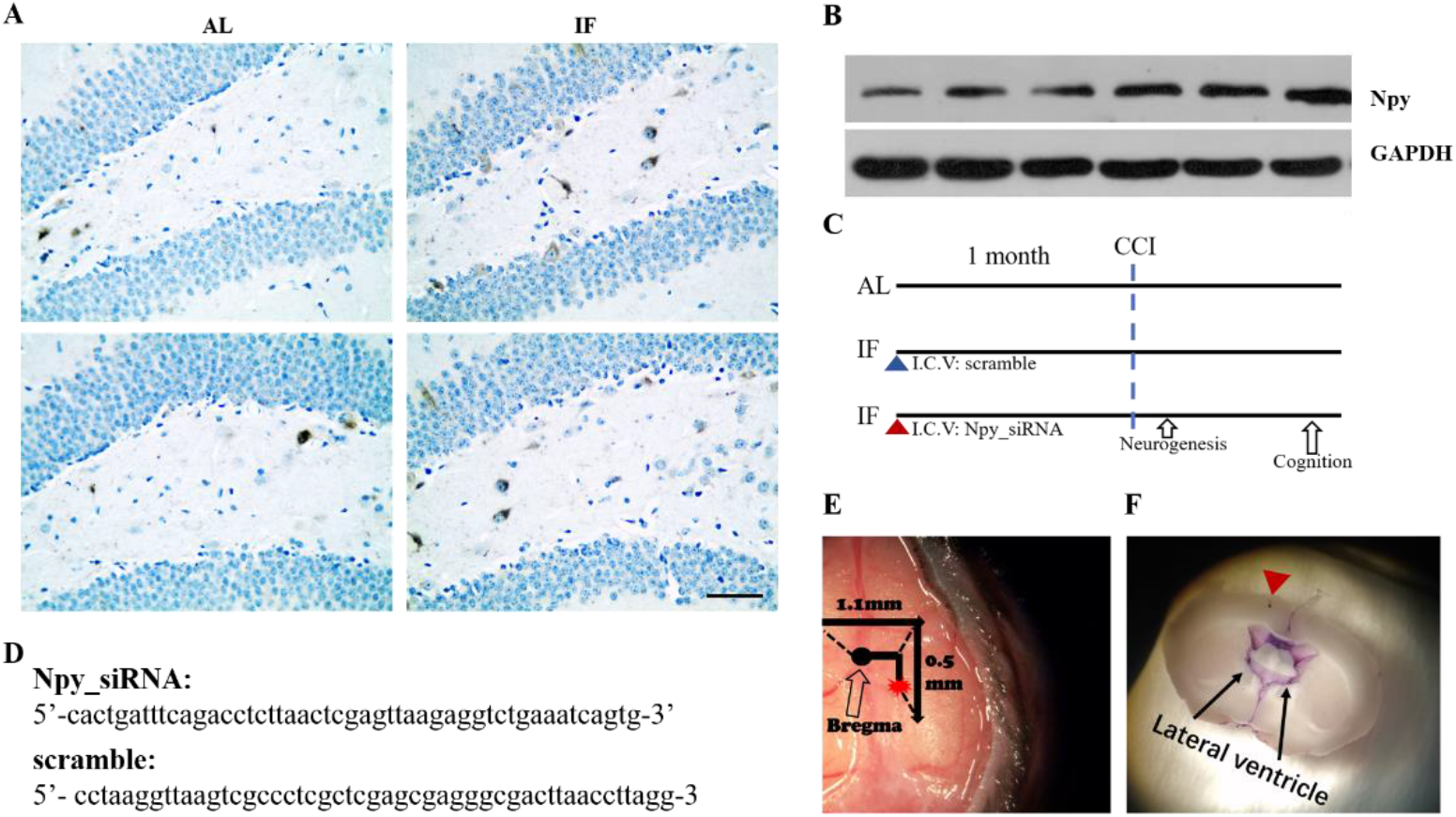
Increased Npy expression may be responsible for the IF-enhanced AHN following TBI. A) more Npy immune-positive cells in dentate gyrus were detected under IF pre-conditioning B) WB results validated the elevated hippocampal Npy expression in the IF group. C) schematics showing timelines for interventions in each group, and examinations post injury. D) sequenced of Npy_siRNA and the scramble. E) injection cite in I.C.V. F) validation of success injection as seen by purple dye in the lateral ventricles.

In order to test whether the IF-enhanced neurogenesis following TBI was mediated by Npy, we used siRNA to locally knock down Npy in vivo (Fig3. C, D). AAV-packaged siRNA was delivered through I.C.V (Fig3. E, F), and the knock-down effect of Npy_siRNA was validated by WB of hippocampus (Fig4. A, B). Subsequently, we conducted immune-staining to examine the neurogenesis post TBI in each group. Results showed that the IF-enhanced neurogenesis was attenuated in the Npy knock-down group, while remained intact under scramble_siRNA treatment (Fig4. C). These results suggested that the IF-enhanced neurogenesis following TBI is mediated via Npy.

**Fig4.**
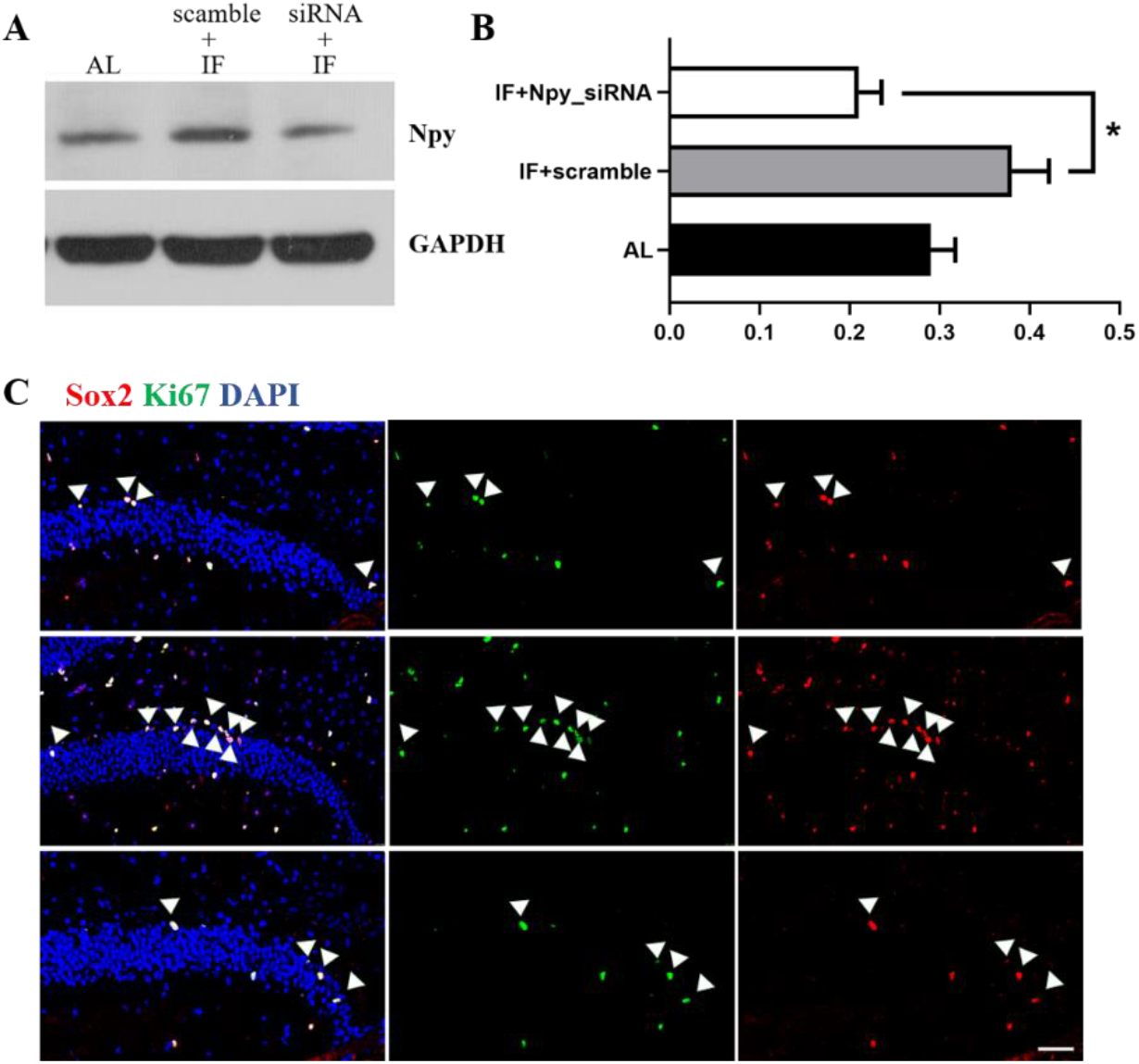
Local knock-down of Npy blocked neurogenesis promoting effect of IF in TBI. A) WB and B) its quantification using ImageJ demonstrated that Npy expression was decreased under siRNA treatment, *P<0.05. C) immunofluorescence results showed that the IF-enhanced neurogenesis was blocked under the Npy knock-down in vivo, scale=100um.

In addition, behavior tests were performed to test spatial learning and memory under Npy_siRNA treatment (Fig5. A-D). We found that the aforementioned neuroprotective effect of IF in cognitive tests was blocked by Npy knock-down (Fig5. A).

**Fig5.**
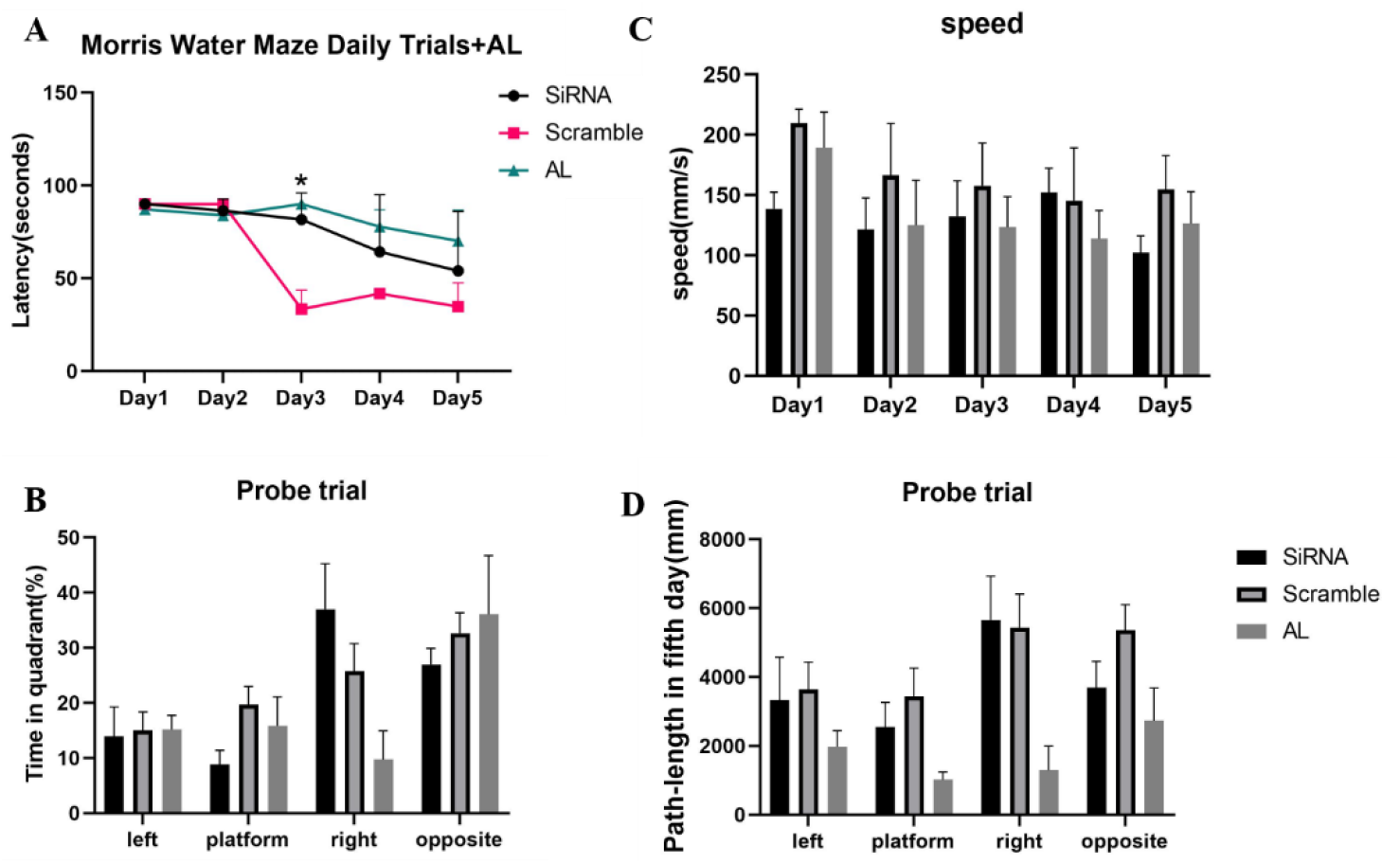
Effect of IF in improving cognitive function post TBI was blocked by Npy knock-down. A) The decreased goal latency in MWM test under IF preconditioning was blocked by in vivo knock-down of Npy (P<0.05). B) time in quadrants C) path length and D) speed tests did not show significant differences among each group.

In conclusion, our results suggested that the neuroprotective role of IF in TBI was mediated by the increase of hippocampal Npy expression.

## Discussion

Over decades, hundreds of rodent studies and clinical trials have revealed the health benefits of fasting in multiple aspects, not simply the results of weight loss[23]. For instance, earlier data has shown that food restriction over a lifetime could prolong lifespan and delay senescence[24]. Subsequently, increasing efforts on the studies in multiple fields revealed positive effects of fasting. Evidences have consistently shown that robust efficacy of caloric restriction in cancers, neurodegenerative disease, epilepsy, ischemic injury, cardiovascular disease, ageing, obesity and its related chronic disorders, etc [25–28]. Among different fasting regimens, the IF was intensely studied recently. Except for the beneficial ketone bodies generated during fasting period, IF induces evolutionarily conserved, adaptive responses that enhance disease resistance and promote self-repairment. During the periodic feeding, tissue specific plasticity or growth is engaged. Given the great benefits of IF as well as the absence of effective interventions for preventing the cognitive defects happened post TBI, we tested the potential therapeutic benefits of IF in the hippocampal behavior and its associated cognitive performance based on a CCI model.

In the present study, we suggest a one-month IF regimen before moderate TBI remarkably ameliorates cognitive dysfunction in the MWM test, 3 months post injury, which has been not been reported previously. As a wealth of evidences have demonstrated that the AHN plays a critical role in the hippocampus related cognition, we then examined the post-injury AHN under different dietary strategies. Consistently, immuno-staining against NSC marker and proliferation marker uncovered enhanced proliferation of NSCs in the SGZ neurogenetic niche under IF. Although a recent study has suggested the relation between IF and AHN[29], their study did not include brain injury. As the microenvironment largely changes under injury condition, the impact study of IF in TBI is still necessary and valuable. In addition, the IF strategy Sang-Ha et al. used entails a 3 months regimen[29], while our study found that a 1 month IF would be able to increase the post-injury AHN and improve the cognitive performance. Furthermore, an animal study suggested that a single 24 hr fasting could bring benefits in TBI, including the increase of mitochondrial oxidative marker and alleviating oxidative stress and calcium loading[30], while our study failed to detect an AHN promoting effect of IF when the mice were given a shorter time of IF regimen (data not shown). To clarify the mechanisms underlying the effect of IF in TBI, we focused on Npy, which has been reported to be involved in feeding and neuro-protecting, respectively[31, 32]. It is wildly known that Npy neurons in the hypothalamus is critical for energy balance[21, 31]. During fasting, hypothalamic Npy expression is largely elevated and thus promotes food consumption and inhibit satiety, and vice versa[31]. Whereas study on the dynamics of hippocampal Npy in the feeding and fasting cycle is quite limited. Hence, we examined the Npy level in the hippocampus responding to fasting and also found an increasement. In previous studies, Npy was suggested to participate in a wide range of events, such as cognitive behavior, epilepsy, chronic stress, depression, social disorders, etc[33, 34]. Furthermore, it has been demonstrated to be a major regulator for the cell proliferation in SVZ-another NSC niche in adult[35]. We validated Npy as an effector of the IF-enhanced AHN and cognitive recovery post TBI by the incomplete loss-of-function study. We delivered Npy-siRNA via I.C.V to locally knock down Npy in vivo and found a blockage of the effects of IF in TBI, eventually help to elucidate the mechanisms underlying the role of IF in TBI. In conclusion, our study suggests that a 1 month IF regime could increase post-injury AHN and alleviate cognitive dysfunction by enhancing hippocampal Npy expression, which may provide new interventions for preventing cognitive impairment after TBI.

